# Interpretable Deep Temporal Structure Learning Model for Early Detection of Alzheimer’s Disease

**DOI:** 10.1101/2019.12.12.874784

**Authors:** Xiaoqian Wang, Dinggang Shen, Heng Huang

## Abstract

In Alzheimer’s research, Mild Cognitive Impairment (MCI) is an important intermediate stage between normal aging and Alzheimer’s disease. How to distinguish MCI samples that finally convert to AD from those do not is an essential problem in the prevention and diagnosis of Alzheimer’s. Traditional methods use various classification models to distinguish MCI converters from non-converters, while the performance is usually limited by the small number of available data. Moreover, previous methods only use the data at baseline time for training but ignore the longitudinal information at other time points along the disease progression. To tackle with these problems, we propose a novel deep learning framework that uncovers the temporal correlation structure of the longitudinal neuroimaing data in the disease progression. In the meantime, we formulate our new deep learning model in an interpretable style such that it provides insights on the important features Alzheimer’s research. We conduct extensive experiments on the ADNI cohort and outperform the related methods with significant margin.

## 1 Introduction

Alzheimer’s disease (AD) is a complex chronic progressive neurodegenerative disease that gradually affects human memory, judgment, and behavior. As an important intermediate stage between normal aging and AD, MCI possesses an increased risk of transiting to AD [31]. That being the case, how to recognize the MCI samples with high potential of switching to AD prior to dementia becomes an essential problem in Alzheimer’s prophylaxis and early treatment.

Neuroimaging provides an effective tool to characterize the structure and functionality of nervous system, thus has greatly contributed to Alzheimer’s study [48]. Extensive work has been proposed to predict MCI conversion using neuroimaging data [34, 17]. Previous methods usually formulate MCI conversion prediction as a binary classification (distinguishing MCI converters from non-converters) [34, 47] or multi-class classification problem (when considering other classes such as AD or health control (HC)) [17, 44], where the methods take the neuroimaging data at baseline time as the input and classify if the MCI samples will convert to AD in years.

Despite the prosperity and progress achieved in MCI conversion prediction, there are still several problems existing in previous methods. 1) Although we expect the model to be capable of forecasting the MCI conversion years before the change of disease status, the training process should not be limited to just baseline data. In the longitudinal study of AD, usually the data at several time points along the disease progression is available, such as baseline, month 6, month 12, *etc*. However, previous methods only consider the baseline data in the training process, thus ignore the temporal correlation structure among other time points. 2) The labeling process for Alzheimer’s is time-consuming and expensive, so the MCI conversion prediction suffers greatly from limited training data.

To deal with these problems, we propose a novel model for MCI conversion prediction. Firstly, we study the temporal correlation structure among the longitudinal data in Alzheimer’s progression. Since AD is a chronically progressive disorder and the neuroimaging features are correlated [27], it can be helpful to analyze the temporal correlation between neuroimaging data in the disease progression as in other nervous system diseases [12]. We construct a regression model to discover such temporal correlation structure between adjacent time points. Our model incorporates the data at all time points along the disease progression and uncovers the variation trend that benefits MCI conversion prediction.

Secondly, we construct a classification model to predict the disease status at each time point. Different from previous classification models that use the baseline data to forecast the progression trend in two or three years, our classification model focuses on adjacent time points. Compared with previous models that require a highly distinguishable conversion pattern appears several years before dementia, our model predicts the progression trend for consecutive time points, thus is more accurate and reliable.

Thirdly, we construct a deep generative model based on generative adversarial network (GAN) to produce more auxiliary data to improve the training of regression and classification model. GAN model is proposed in [15], which uses the adversarial mechanism to learn the inherent data distribution and generate realistic data. We use the generative model to learn the joint distribution of neuroimaging data at consecutive time points, such that more reliable training data can be obtained to improve the prediction of MCI conversion.

Last but not least, we propose a new idea to improve the interpretability of our model. Interpretability of the model is very important to inspect the validity of the prediction. In medical diagnosis, it is crucial to verify that the diagnosis is based on valid reasons. A thorough knowledge of the model behavior is essential to the enhance end-user’s understanding and trust in the model. Moreover, good interpretability of the model can improve the performance of the model [2], strengthen the explanation of an algorithm’s decision[16, 20], and enable the discovery of new science [36]. In this paper, we formulate our deep learning model in a novel interpretable manner in order to improve the understanding of the predictive mechanism and provide insights on important features in Alzheimer’s disease research.

## 2 Related Work

Here we review related works from three perspectives: MCI conversion prediction, constructing an interpretable model, and making interpretation for a given black-box model.

### 2.1 MCI Conversion Prediction

Recent advances in machine learning have enabled several new models for MCI conversion prediction. In [17], linear SVM method was employed to distinguish between MCI converters and non-converters, where the extraction of multi-scale features from baseline MRI data strengthened the classification. In [47], the authors used a non-linear classification model to flexibly describe the complex relationship between neuroimaging data and disease status. In [44], the authors integrated more information from health control and patient samples in MCI conversion prediction with a multi-class classification model and automatically uncovered the interrelations among different classes to improve the classification performance. The widespread application of machine learning models in MCI conversion prediction enables better understanding of the progression patterns in MCI, thereby contributing to the prevention and diagnosis of Alzheimer’s.

### 2.2 Interpretable Model Construction

There are several recent work on building interpretable models using sparse linear models, decision trees and rules lists owing to the intrinsic interpretable property of these methods. These models are very useful when interpretation of the computational model is important. For example, [43] propose a sparse linear model to analyze Alzheimer’s disease related brain regions. [24] use the decision sets to construct an interpretable classifier for stroke prediction. [26] use rules list to build an interpretable model for stroke prediction while [41] adopt rules list to predict hospital readmissions. However, the performance of these interpretable models are limited by the complexity constraints. The number of non-zeros coefficients in sparse linear models, and the number and length of rules in rules lists and decision sets have to be carefully constrained, such that the model is not too complex to interpret. Such constrains limit the flexibility of the interpretable models, making it difficult to apply to large-scale and complex tasks.

### 2.3 Interpretation of a Black-Box Model

Recent methods on interpreting a black-box model can be roughly divided into three categories: feature visualization, attribution and model approximation. In feature visualization [11, 13, 30], the interpretation is based on finding the input that activates a certain output. From the simulated input, human users can analyze the model’s understanding in the data by visualizing the desired input of the model. Moreover, attribution methods [4, 29, 49, 39, 37, 36, 3] propose to identify which part of input is the most responsible for the output by tracking from the output backward to the input. Attribution methods show the role of each feature and help the users to interpret the contribution of the features in the prediction. Model approximation is the method that builds a simpler interpretable model to approximate the black-box model and use the explanation from the simpler model for explanation. In [32], Ribeiro *et al*. build a sparse linear model to approximate any given classifier such that the output of the linear model is locally consistent to the output from the black-box classifier. In [5], Bastani *et al*. use decision trees to approximate the black-box model for a global interpretation of the model behavior. Also, there are other model interpretation methods in addition to the above three categories. [23] formulates the interpretation by analyzing the impact of each training sample to the model and picking out the most influential training samples in the prediction.

Although the recent works have introduced advances in model interpretation and strengthened the understanding in black-box models, there still lacks a universal standard in evaluating and quantifying the quality of interpretation [10]. [21] points out that several well-known interpretation methods, including DeConvNet [49], Guided BackProp [39] and LRP [4], fail to provide theoretically correct interpretation for a simple linear model. In [1], Adebayo *et al*. points out that a deep neural network (DNN) with random weights shares both visual and quantitative similarity with a DNN with learned weights on the learned interpretation. All of these findings raise concerns about the quality and consistency of interpretation. In [8], Chu *et al*. propose a theoretical analysis on the consistency of the interpretation, whose method is, however, constrained to piecewise linear models and is not applicable to neural networks with other standard layers like batch normalization, *etc*.

It is notable that our interpretable temporal structure learning model is different from the previous works from several different aspects. In terms of MCI conversion prediction, we propose a novel idea of uncovering the progressive trend of Alzheimer’s disease to improve the prediction of MCI conversion status. Moreover, unlike previous models that predict the MCI conversion two or three years before the disease status changes, our prediction focuses on adjacent time points thus is more accurate.

In terms of model interpretation, our model differs from the model interpretation methods in that we directly construct an interpretable model, guaranteeing that the interpretation is consistent with the model behavior. In addition, different from previous interpretable models whose performance is limited by the interpretable constraints, our model is flexible and achieves roughly comparable performance with state-of-the-art black-box structures on difficult tasks such as image classification and sentence classification.

## 3 Constructing an Interpretable Deep Learning Model in An Additive Manner

In this section, we first introduce how to build a deep learning model in an interpretable manner such that the model provides direct explanation on what features play an important role in the prediction. In a supervised learning problem, suppose the data is drawn from the distribution *p*(**x**, **y**), traditional black-box models propose to find a prediction function *f* that optimizes the following:

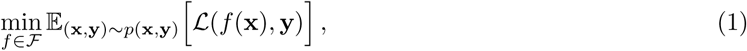

where ℒ is the loss function and ℱ is the assumptions and constraints on the structure of *f*. In deep learning models, function *f* is non-linear and complicated, which makes the behavior of function *f* difficult to understand. For a given data **x**, such black-box structure renders the mechanism behind how the model makes the prediction **ŷ** = *f* (**x**) to be opaque and not interpretable.

One method to understand the behavior of *f* is to find an interpretable model *h* (*e*.*g*, sparse linear model *h*(**x**) = **x**^T^*W*) to locally approximate *f* [32]. It is notable that the coefficient matrix *W* is dependent on the data **x** to be interpreted. As a consequence, we can define *W* as a function *w*.*r*.*t*. the data **x** as *W* = *g*(**x**) and formulate the prediction as **ŷ** = **x**^T^*g*(**x**). We plug **ŷ** = **x**^T^*g*(**x**) in Eq. (1) and propose to optimize the following:

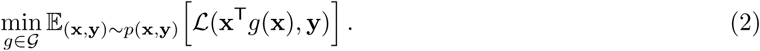

where *𝒢* is the assumptions and constraints on the structure of function *g*. Note that in Problem (2), *g*(**x**) makes an explicit interpretation on the contribution of features in **x** to **ŷ**, such that the mechanism behind the model behavior is clear.

Given a set of data samples 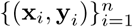, we optimize the following empirical loss to approximate the expected loss in Problem (2):

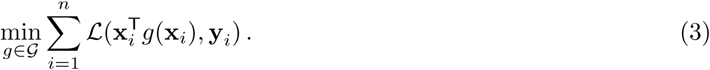

For the sake of interpretability, we impose sparsity constraints on the learned coefficient matrix to limit the number of non-zero values in the coefficient matrix for each data and propose the following objective function for our method:

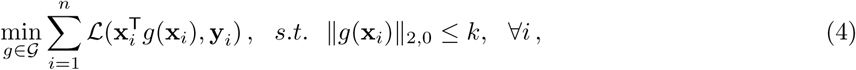

where *k* is the number of non-zeros values in each row of the coefficient matrix.

We would like to point out that the formulation in Problem (4) enjoys several advantages:

– The model is clearly interpretable since the coefficient matrix *g*(**x**_*i*_) makes an explicit and direct explanation on the association between data **x**_*i*_ and the prediction 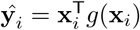.
– The function *g* enjoys the representative power and flexibility of the black-box models, thus our model can perform well in the cases when *f* works well in Problem (1).
– Our interpretable learning idea is not restricted to any specific structure and can adapt to many state-of-the-art architecture.

For simplicity, we plot an illustration figure in Fig. 2 to explain our idea of building an interpretable model. The learned coefficient matrix on MNIST data for digit classification indicates that our model correctly classifies the digit according to reasonable features, showing that our model is both effective and trustworthy.

**Fig. 1.**
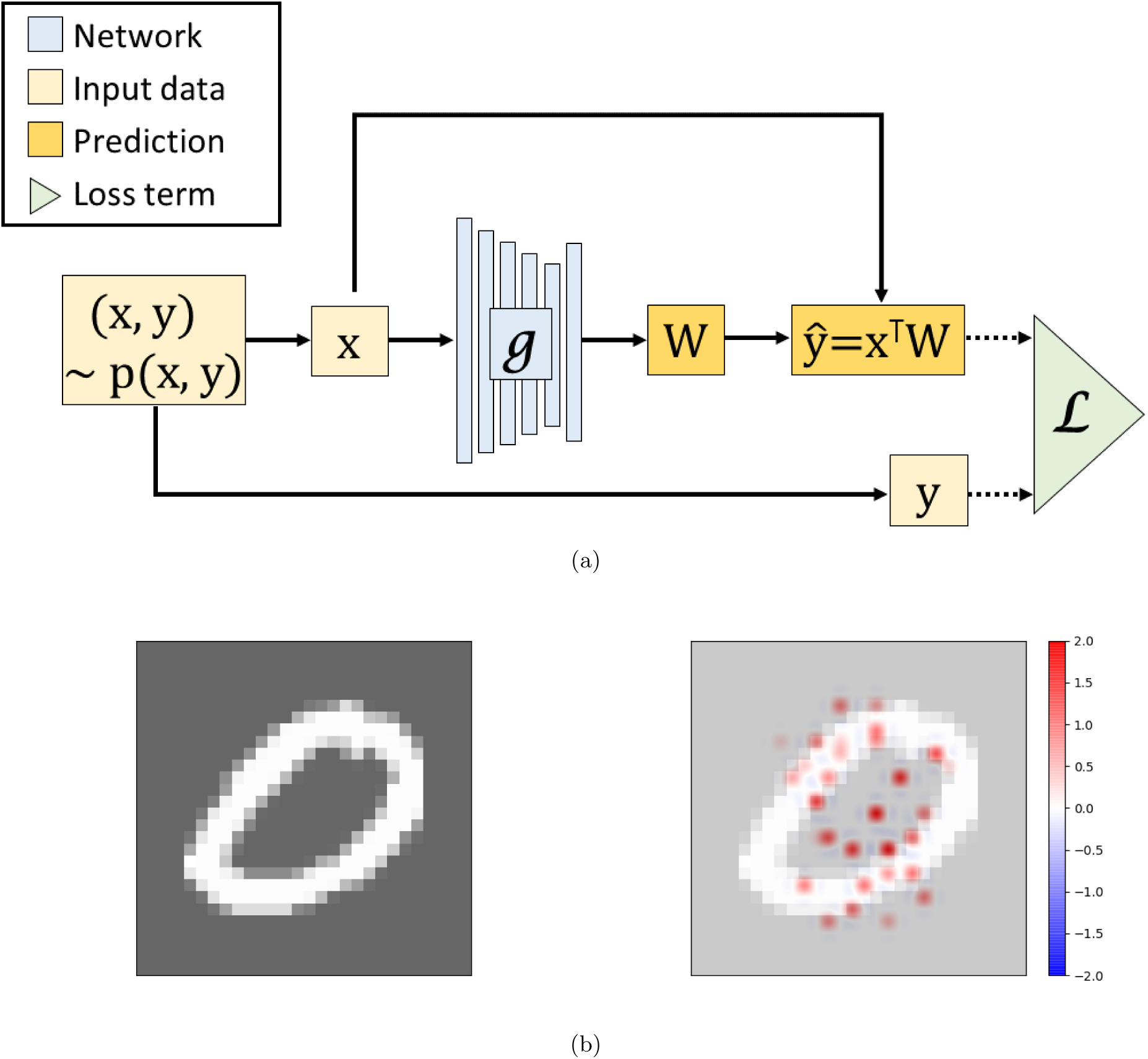
Illustration of our model. (a) Structure of our idea of formulating the model in an interpretable way. Different from previous black-box models that learn the mapping from the input **x** to the response variable **y**, our model learns the coefficient matrix *W* for a given **x** and use **x**^T^*W* as the prediction. (b) A data example in the MNIST dataset and illustration of the corresponding coefficients learned from our model.

**Fig. 2.**
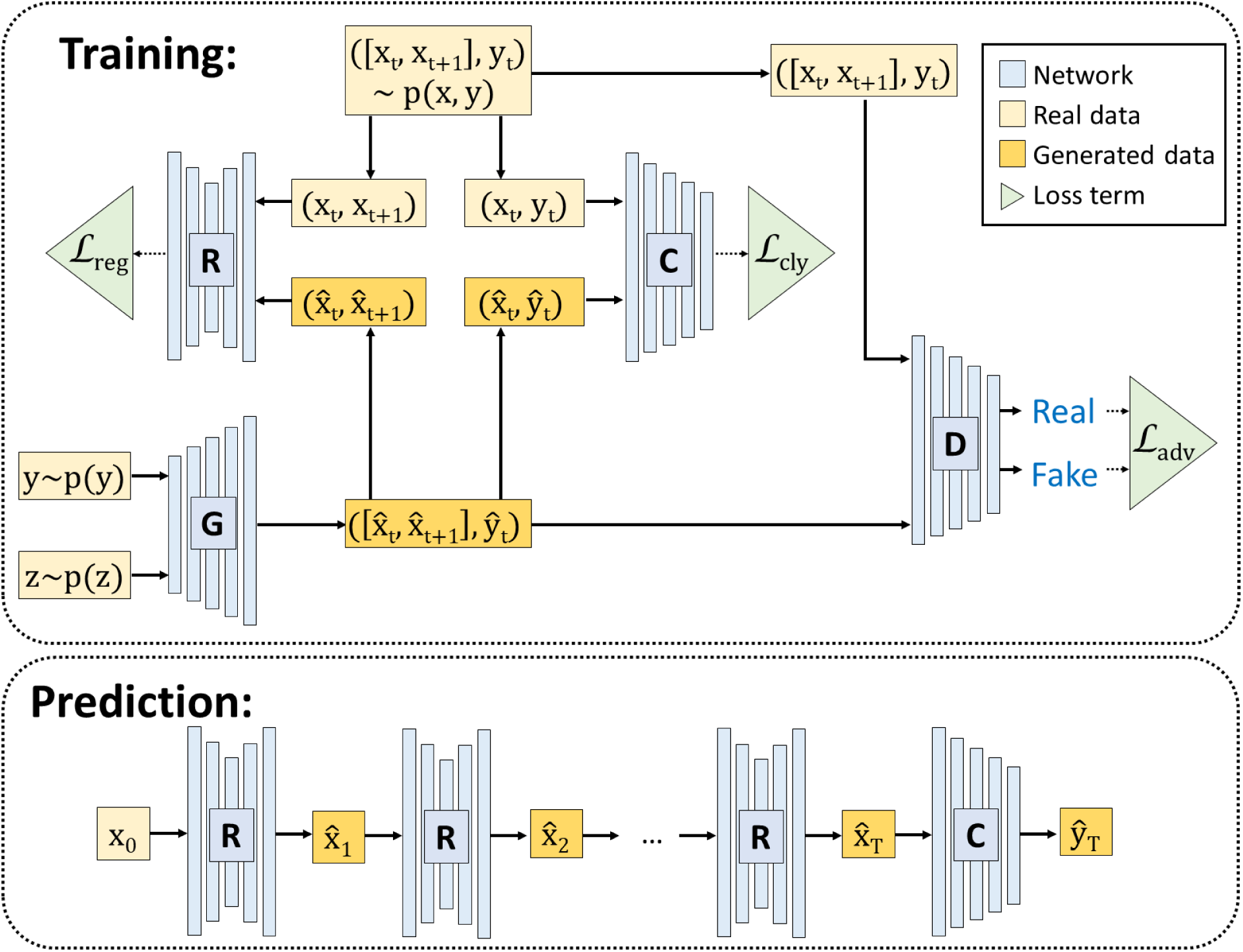
Illustration of our ITGAN model. **x**_*t*_ and **x**_*t*+1_ are the neuroimaging data at two adjacent time points and **y**_*t*_ is the label (**y**_*t*_ = 1 if both **x**_*t*_ and **x**_*t*+1_ are at AD status; **y**_*t*_ = 2 if **x**_*t*_ is MCI while **x**_*t*+1_ is AD; **y**_*t*_ = 3 if both **x**_*t*_ and **x**_*t*+1_ are MCI.). The regression network *R* predicts **x**_*t*+1_ from **x**_*t*_ so as to uncover the temporal correlation between adjacent time points. The classification network *C* predicts the label **y**_*t*_ from **x**_*t*_. We also construct a generative model with generator *G* and discriminator *D* to approximate the joint distribution underlying data pair ([**x**_*t*_, **x**_*t*+1_], **y**_*t*_) to generate more reliable data for training *R* and *C*. In the prediction process, the neuroimaging data **x**_0_ at baseline time for MCI samples is given, and we use *R* and *C* to predict whether the MCI sample will convert to AD at time *T*. It is notable that our *C* model is constructed in an interpretable manner as illustrated in Section 3, such that we can get direct interpretation on the important features used in the prediction.

## 4 Temporal Correlation Structure Learning Model

In this section, we introduce our idea of learning the temporal correlation structure in the longitudinal neuroimaging data for MCI conversion prediction.

### 4.1 Problem Definition

In MCI conversion prediction, for a certain sample and a time point *t*, we use **x**_*t*_ ∈ ℝ^*p*^ to denote the neuroimaging data at time *t* while **x**_*t*+1_ ∈ ℝ^*p*^ for the next time point, where *p* is the number of imaging markers. **y**_*t*_ ∈ ℝ is the label showing the disease status at time *t* and *t* + 1. Here we define three different classes for **y**_*t*_: **y**_*t*_ = 1 means the sample is AD at both time *t* and *t* + 1; **y**_*t*_ = 2 shows MCI at time *t* while AD at time *t* + 1; while **y**_*t*_ = 3 indicates that the sample is MCI at both time *t* and *t* + 1. In the prediction, given the baseline data of an MCI sample, the goal is to predict whether the MCI sample will finally convert to AD or not.

### 4.2 Revisit GAN Model

GAN model is proposed in [15], which plays an adversarial game between the generator *G* and discriminator *D*. The generator *G* takes a random variable **z** as the input and outputs the generated data to approximates the inherent data distribution. The discriminator *D* is proposed to distinguish the data **x** from the real distribution and the data produced from the generator. Whereas the generator *G* is optimized to generate data as realistic as possible to fool the discriminator. The objective function of the GAN model has the following form.

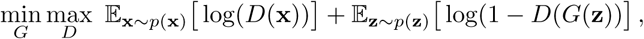

where *p*(**z**) denotes the distribution of the random variable and *p*(**x**) represents the distribution of real data. The min-max game played between *G* and *D* improves the learning of both the generator and discriminator, such that the model can learn the inherent data distribution and generate realistic data.

### 4.3 Illustration of Our Model

Inspired by [7], we propose to approximate the joint distribution of neuroimaging data at consecutive time points and data label ([**x**_*t*_, **x**_*t*+1_], **y**_*t*_) ∼ *p*(**x**, **y**) by considering the following:

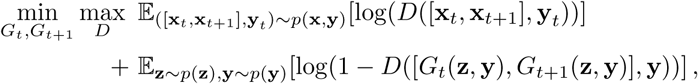

where the generators take a random variable **z** and a pseudo label **y** as the input and output a data pair ([*G*_*t*_(**z**, **y**), *G*_*t*+1_(**z**, **y**)], **y**) that is as realistic as possible. Still, the discriminator is optimized to distinguish real from fake data. The construction of such generative model approximates the inherent joint distribution of neuroimaging data at adjacent time points and label, which generates more reliable samples for the training process.

To uncover the temporal correlation structure among the neuroimaging data between consecutive time points, we construct a regression network *R* to predict **x**_*t*+1_ from **x**_*t*_, such that progression trend among neuroimaging data along the disease progression can be learned. The network *R* takes data from both real distribution and the generators as the input and optimize the following:

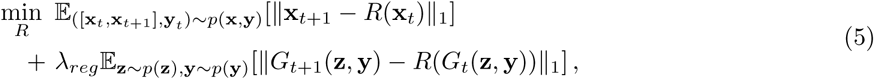

where the hyper-parameter *λ*_*reg*_ balances the importance of real and generated data. We consider 𝓁_1_-norm loss to make the model *R* more robust to outliers.

In addition, we construct a classification structure *C* to predict the label **y**_*t*_ given data **x**_*t*_. The optimization of *C* is based on the following:

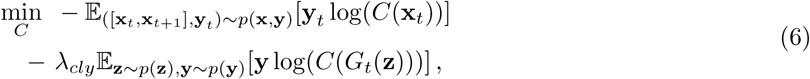

where *λ*_*cly*_ is a hyper-parameter to balance the role of real and generated data.

Given a set of real data 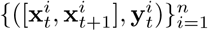, the above three loss terms can be approximated by the following empirical loss:

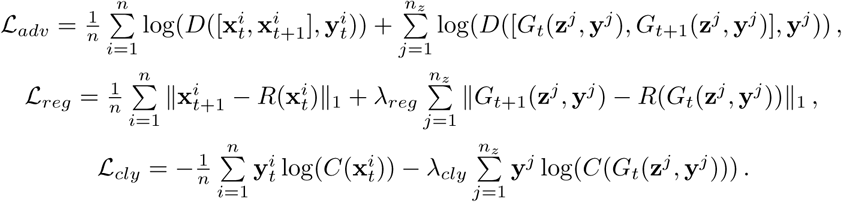

For a clear illustration, we plot a figure in Fig. 2 to show the structure of our ITGAN model (interpretable temporal correlation structure learning for MCI conversion prediction with GAN). The implement details of the networks can be found in the experimental setting section. The optimization of our model is based on a variant of mini-batch stochastic gradient descent method.

## 5 Experimental Results

### 5.1 Experimental Setting

To evaluate our ITGAN model, we compare with the following methods: **SVM-Linear**(support vector machine with linear kernel), which has been widely applied in MCI conversion prediction [17, 34]; **SVM-RBF** (SVM with RBF kernel), as employed in[25, 47]; and **SVM-Polynomial** (SVM with polynomial kernel) as used in [25]. Also, to validate the improvement by learning the temporal correlation structure, we compare with the **Neural Network** with exactly the same structure in our classification network (network *C* in Fig. 2) that only uses baseline data. Besides, we compare with the case where we do not use the GAN model to generate more auxiliary samples, *i*.*e*., only using network *C* and *R* in Fig. 2, which we call **Temporal-Deep**.

The classification accuracy is used as the evaluation metric. We divide the data into three sets: training data for training the models, validation data for tuning hyper-parameters, and testing data for reporting the results. We tune the hyper-parameter *C* of SVM-linear, SVM-RBF and SVM-Polynomial methods in the range of {10^−3^, 10^−2^, …, 10^3^}. We compare the methods when using different portion of testing samples and report the average performance in five repetitions of random data division. It is notable that our classification network *C* output the weight matrices *W* such that the prediction of *C* is intrinsically interpretable. We plot an illustration figure in Fig. 3.

**Fig. 3.**
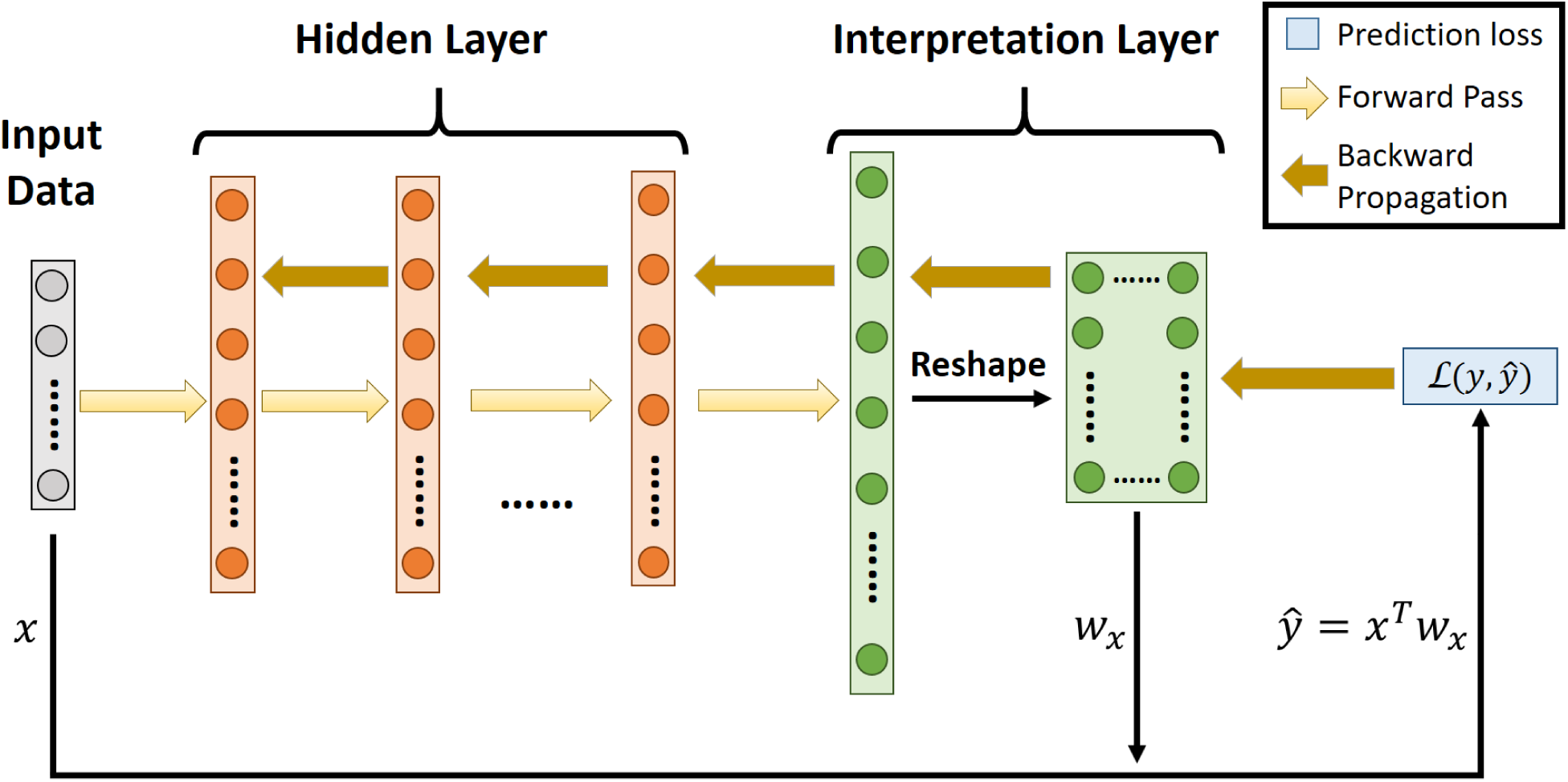
Illustration of our idea on formulating a deep learning model in an interpretable manner.

In our ITGAN model, we use the fully connected neural network structure for all the networks *G, D, R* and *C*. The implementation detail is as follows: The number of each hidden layer contains 100 hidden units while the number of hidden layers in the structure *G, D, R* and *C* is 2, 1, 3, 4 respectively. We use leaky rectified linear unit (LReLU) [28] with leakiness ratio 0.2 as the activation function of all layers except the last layer and consider weight normalization [33] for layer normalization. We adopt the dropout mechanism for all layers except the last layer with the dropout rate of 0.1 [40]. The weight parameters of all layers are initialized using the Xavier approach [14]. We use the ADAM algorithm [22] to update the weight parameters with the hyper-parameters of ADAM algorithm set as default. Both values of *λ*_*reg*_ in Eq. (5) and *λ*_*cly*_ in Eq. (6) are set as 1.0. We use the generator *G* to produce ten times of the samples from the real distribution. We use Theano toolbox for the implementation. All experiments are conducted on a machine with one Titan X pascal GPU.

### 5.2 Data Description

Data used in this paper was downloaded from the ADNI database (adni.loni.usc.edu). Each MRI T1-weighted image was first anterior commissure (AC) posterior commissure (PC) corrected using MIPAV2, intensity inhomogeneity corrected using the N3 algorithm [38], skull stripped [46] with manual editing, and cerebellum-removed [45]. We then used FAST [50] in the FSL package3 to segment the image into gray matter (GM), white matter (WM), and cerebrospinal fluid (CSF), and used HAMMER [35] to register the images to a common space. GM volumes obtained from 93 ROIs defined in [18], normalized by the total intracranial volume, were extracted as features. Out of the 93 ROIs, 24 disease-related ROIs were involved in the MCI prediction, where the selection of AD-related ROIs is based on [42]. This experiment includes data from six different time points: baseline (BL), month 6 (M6), month 12 (M12), month 18 (M18), month 24 (M24) and month 36 (M36). All 216 samples with no missing MRI features at BL and M36 time are used by all the comparing methods, where there are 101 MCI converters (MCI at BL time while AD at M36) as well as 115 non-converters (MCI at both BL and M36). Since our ITGAN model can use data at time points other than BL and M36, we include a total of 1419 data pairs with no missing neuroimaging measurement for training the classification, regression and generative model in our ITGAN model. All neuroimaging features in the data are normalized to zero mean and unit variance.

### 5.3 MCI Conversion Prediction

We summarize the MCI conversion classification results in Table 5.3. The goal of the experiment is to accurately distinguish converter subjects from non-converters among the MCI samples at baseline time. From the comparison we notice that ITGAN outperforms all other methods under all settings, which confirms the effectiveness of our model. Compared with SVM-Linear, SVM-RBF, SVM-Polynomial and Neural Network, the ITGAN and Temporal-Deep model illustrates apparent superiority, which validates that the temporal correlation structure learned in our model substantially improves the prediction of MCI conversion. The training process of our model takes advantage of all the available data along the progression of the disease, which provides more beneficial information for the prediction of MCI conversion. By comparing ITGAN and Temporal-Deep, we can notice that ITGAN always performs better than Temporal-Deep, which indicates that the generative structure in ITGAN could provide reliable auxiliary samples to strengthen the training of regression *R* and classification *C* model, thus improves the prediction of MCI conversion.

**Table 1.** MCI conversion prediction with different portion of testing data. This table shows the average classification accuracy and standard deviation in 5 repetitions. The best results are marked in bold.

| Methods | 10% | 20% | 50% |
| --- | --- | --- | --- |
| SVM-Linear | 0.6273±0.0668 | 0.6558±0.0648 | 0.6093±0.0484 |
| SVM-RBF | 0.5818±0.0881 | 0.5767±0.0770 | 0.5852±0.0381 |
| SVM-Polynomial | 0.6545±0.0617 | 0.5953±0.1253 | 0.5611±0.0429 |
| Neural Network | 0.3727±0.0340 | 0.4233±0.0426 | 0.4685±0.0324 |
| Temporal-Deep | 0.7455±0.0680 | 0.7442±0.0416 | 0.7556±0.0359 |
| ITGAN | **0.8091±0.0881** | **0.7488±0.0271** | **0.7611±0.0301** |

**Table 2.** Classification accuracy comparison when involving all 93 ROIs in MCI conversion prediction.

| Methods | 10% | 20% | 50% |
| --- | --- | --- | --- |
| SVM-Linear | 0.6182±0.1206 | 0.6140±0.0315 | 0.5815±0.0515 |
| SVM-RBF | 0.6727±0.1164 | 0.5581±0.0510 | 0.6056±0.0404 |
| SVM-Polynomial | 0.6182±0.1616 | 0.6558±0.1034 | 0.5981±0.0471 |
| Neural Network | 0.4455±0.1052 | 0.4605±0.0451 | 0.4593±0.0126 |
| Temporal-Deep | 0.7727±0.0498 | 0.7814±0.0479 | 0.7000±0.0313 |
| ITGAN | **0.8091±0.0668** | **0.7860±0.0372** | **0.7148±0.0416** |

### 5.4 Visualization of the Imaging markers

In Fig. 4, we illustrate the brain map corresponding to the top 15 imaging features that are learned in our *C* model. In this subsection, we use all 93 ROIs in the MCI conversion prediction and validate if the top neuroimaging features captured by our ITGAN model is Alzheimer’s disease relevant.

**Fig. 4.**
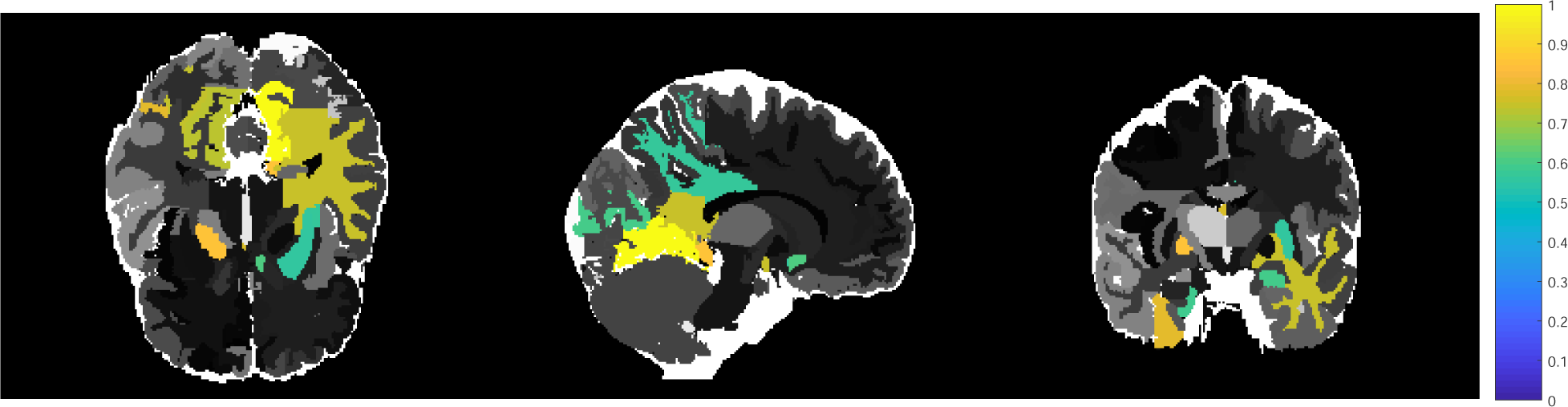

Among the detected neuroimaging features, we identified several Alzheimer’s relevant regions of interests (ROIs) that have been replicated in previous literature. In our classification *C* model, medial occipitotemporal gyrus left and inferior temporal gyrus right rank play an important role in the early detection of MCI. Convit *et al*. [9] ascertained the influence of the atrophy in the medial occipitotemporal, inferior, and middle temporal gyri in the decline to Alzheimer’s disease. Moreover, our ITGAN model also validated temporal lobe WM left as a top feature to predict the conversion of MCI, the this finding has been replicated in several previous works [19, 6]. The replication of this findings validated the effectiveness of our interpretation method, such that our ITGAN model provides direct explanation of the important features for predicting the conversion pattern of MCI subjects, and the explanation coincides with previous research on the important features for MCI subjects declining to Alzheimer’s disease.

### 6 Conclusion

In this paper, we proposed a novel ITGAN model for MCI conversion prediction. Our model considered the data at all time points along the disease progression and uncovered the temporal correlation structure among the neuroimaing data at adjacent time points. We constructed a generative model to produce more reliable data to strengthen the training process. Moreover, our ITGAN model is constructed in an interpretable manner such that we can get direct understanding of the important neuroimaging features in the prediction. Our model illustrated superiority in the experiments on the ADNI data.

